# The Impact of Chronic Phthalate Exposure on Rodent Anxiety and Cognition

**DOI:** 10.1101/2023.04.13.536567

**Authors:** Zhe Yu, Laxmi Iyer, Adam P. Swiercz, Elizabeth Paronett, Manelle Ramadan, Paul J. Marvar, Nikki Gillum Posnack

**Affiliations:** Department of Pharmacology and Physiology, George Washington University, Washington DC; NIMH, National Institutes of Health, Bethesda, MD; Children's National Heart Institute, Sheikh Zayed Institute for Pediatric Surgical Innovation, Children's National Hospital, Washington, DC; Department of Psychiatry and Behavioral Sciences, George Washington University, Washington DC; Uniformed Services University Health Sciences

**Keywords:** Phthalate, Anxiety, Cognition, Corticosterone

## Abstract

There is a growing importance for environmental contributions to psychiatric disorders and understanding the impact of the exposome (i.e., pollutants and toxins). Increased biomonitoring and epidemiological studies, for example, suggest that daily phthalate chemical exposure contribute to neurological and behavioral abnormalities, however these mechanisms remain poorly understood. The current study therefore aimed to examine the effects of chronic phthalate exposure on rodent anxiety behaviors, cognition, and the impact on hypothalamic-pituitary­ adrenal (HPA)-axis function. Adult male mice (C57BL6/J) were administered mono-2-ethylhexyl phthalate (MEHP) via drinking water (1 mg/ml), and anxiety-like behavior, cognition combined with HPA- axis and inflammatory assays were assessed after 3 weeks of MEHP exposure. MEHP-treated mice exhibited enhanced generalized anxiety-like behaviors, as demonstrated by reduced time spent in the open-arm of the elevated plus maze (EPM) and center exploration in the open field (OF). Tests of spatial, cognition and memory function were unchanged. Following MEHP administration, circulating levels of corticosterone and pro­ inflammatory cytokines were significantly increased, while at the tissue level, MEHP-dependent reductions in glucocorticoid metabolism genes 11β-hydroxysteroid dehydrogenase (11β-HSD) 1 and 2. These data suggest that chronic MEHP exposure leads to enhanced generalized-anxiety behaviors independent of rodent measures of cognition and memory, which maybe driven by MEHP-dependent effects on HPA-axis and peripheral glucocorticoid metabolism function.

## Introduction

Phthalates are synthetic chemicals, commonly used as plasticizers to impart flexibility to polyvinyl chloride (PVC) for use in plastic consumer and medical products (Afshari et al., 2004). Despite the inherent benefits of plastics, the relative abundance of phthalate chemicals in the environment has more recently raised concerns pertaining to human health and disease risk (Posnack, 2021; Ramadan et al., 2020). Human exposure to phthalates can occur through chemical leaching or migration, since the phthalate plasticizer is not covalently bound to the PVC matrix. Given the ubiquity of plastic materials in the environment, human biomonitoring studies indicate widespread daily exposure to phthalates, with detectable levels in >75% of the general population (Calafat et al., 2017; Kato et al., 2004; Silva et al., 2004; Zota et al., 2014). Therefore additional studies to elucidate the impact of phthalate exposure on biological pathways, specifically related to the growing concern for the impact of phthalate exposure on brain development and mental health(Cardenas-Iniguez et al., 2022), are needed.

Phthalates are classified as endocrine-disrupting chemicals and growing clinical and pre­ clinical evidence has found that phthalate exposure interferes with neurodevelopment and increases the risk for behavioral disorders (Braun, 2017). For example, following prenatal or early childhood phthalate exposures, there are multiple epidemiological and clinical based studies that have detected associations for adverse cognitive and neurobehavioral outcomes (Ejaredar et al., 2015; Zhang et al., 2019), including somatic symptom disorder (Kobrosly et al., 2014), attention-deficit/hyperactivity disorder-related behaviors (Chopra et al., 2014; Engel et al., 2021; Kim et al., 2009; Shoaff et al., 2020), externalizing behavior including aggression, depression, reduced emotional control (Lien et al., 2015; Shoaff et al., 2019), and anxiety proneness (Park et al., 2015). The causal and/or consequential mechanisms and neurobiological pathways underlying the impact of postnatal phthalate exposure on cognitive and behavioral outcomes, however, remain poorly understood (Carbone et al., 2019).

Using a rodent model, the current study therefore aimed to examine the effects of postnatal chronic phthalate exposure on anxiety, memory, and cognitive and neuroendocrine function. Our results provide new evidence demonstrating that chronic phthalate exposure contributes to enhanced anxiety-like behaviors, which may be driven by phthalate-dependent alterations in neuroendocrine and glucocorticoid metabolism function.

## Materials and Methods

### Animals

All experimental protocols were approved by the Institutional Care and Use Committee of The George Washington University and were in compliance with National Institutes of Health guidelines. Adult male C57BL/6J mice (10 weeks old, n=60) used in this study were from Jackson Laboratory (Bar Harbor, ME, USA), and were housed in a temperature­ and humidity-controlled room on a 12-hour light/dark cycle with water and food available ad libitum.

### MEHP administration

Mice received 1 mg/ml MEHP administration through drinking water. To optimize the aqueous solubility of MEHP, drinking water was also supplemented with 20 mg/ml captisol cyclodextrin (Cydex Pharmaceuticals, Lawrence, KS, USA). Vehicle control animals were similarly administered 20 mg/ml captisol through drinking water. After 3 weeks treatment, mice underwent behavioral testing and were subsequently sacrificed at the end of the 5th week. Two cohorts of mice were used for behavioral testing (**Figure 1**); animals received MEHP or vehicle-control drinking water during the entire testing period.

**Figure 1:**
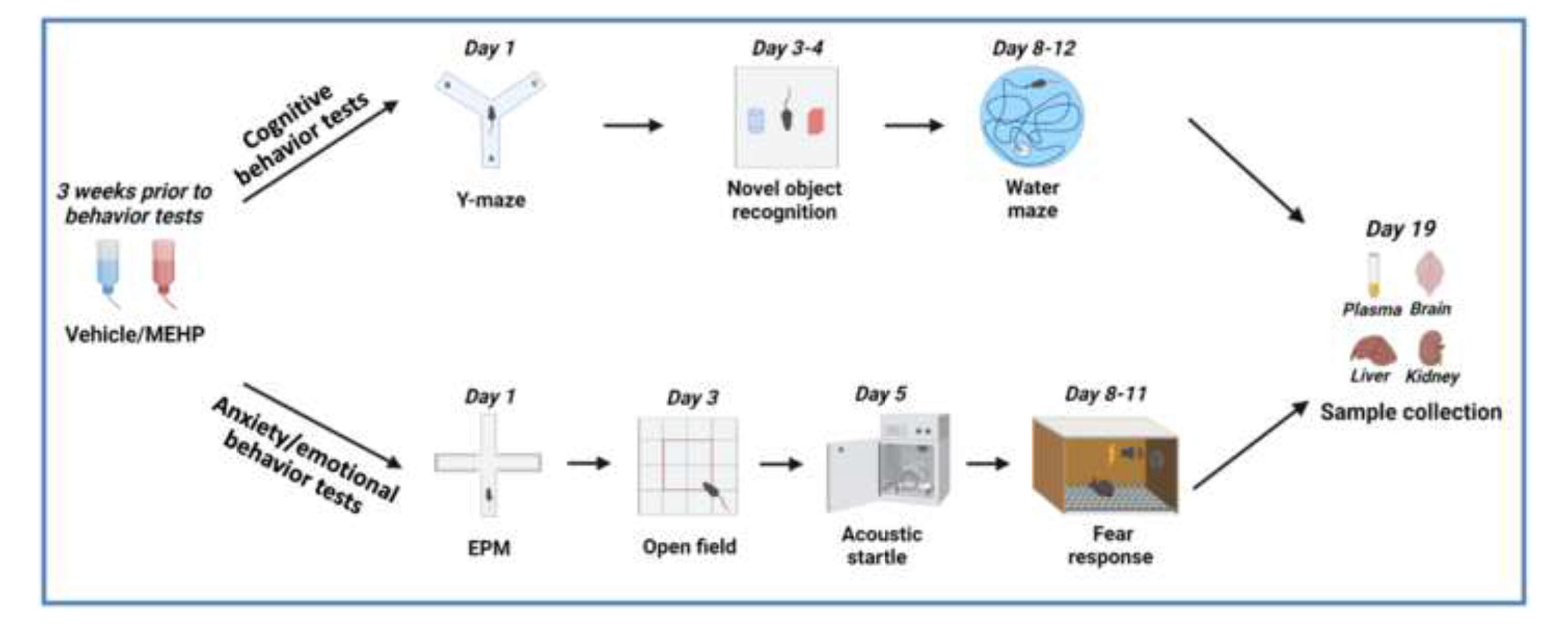
Experimental Protocol and Timeline: Three weeks following MEHP, mice underwent either cognitive behavior tests or anxiety/emotional behavior tests, followed by blood and tissue sample collection. EPM, elevated plus maze.

### Behavioral Testing

To decrease the possible stress caused by room changes, 48 hours prior to the behavior tests, mice were brought to the behavior testing room for testing environment habituation. Animal activities were tracked and analyzed via the ANY-maze video tracking software (Stoelting Co., Wood Dale, IL, USA). The freezing behavior during fear response tests was recorded and quantified using Freezeframe 3.32 (Coulboum Instruments, Holliston, MA, USA). The startle reflex was evaluated by the SR-Lab Startle Systems (San Diego Instruments, San Diego, CA, USA). Additional details are provided in the Supplement.

### Tissue and plasma collection

After behavioral tests were complete, mice were sacrificed via decapitation and the brain, liver, and kidneys were dissected and fresh frozen with liquid nitrogen. Whole blood was collected in EDTA-coated tubes (Thermo Fisher Scientific, Waltham, MA, USA), samples were centrifuged (5000 g, 10 min) and plasma was collected. Tissues and plasma samples were stored at −80 °C for future study.

### Enzyme-linked immunosorbent assay (ELISA)

Plasma corticosterone (AR E-8100, LDN, Nordhom, Germany), adrenaline/noradrenaline/dopamine (BA E-5600, LDN, Nordhom, Germany), adrenocorticotropic hormone (ACTH, M046006, MD Bioproducts, Zurich, Switzerland), and aldosterone (ADI-900-173, LDN, Nordhom, Germany) were measured using commercially available ELISA kits according to the manufacturer’s instructions.

### Cytokine Analysis

Plasma pro-inflammatory cytokines were measured using commercially available electrochemiluminescence kits (Meso Scale Discovery, Gaithersburg, Maryland, USA). For the V-Plex Plus Proinflammatory Panell Mouse Kit (K-15048Gl), the plasma samples were run in duplicate at 1:2 dilution and the assay was performed according to the manufacturer’s instructions.

### RT-qPCR

The hypothalamus was collected by a 1-mm diameter brain tissue punch. Total RNA was extracted from hypothalamus, liver, and kidney tissue using Trizol reagent (Thermo Fisher Scientific, Waltham, MA, USA) according to the manufacturer’s instructions. Additional details can be found in the Supplement.

### Data Presentation and Statistical Analysis

For all data analysis, distributions were checked for non-trivial violation of normality assumptions using graphical methods and checked for outliers using the Grubbs outlier test (a=0.05) or ROUT (Q=l %) in Graphpad Prism 9.0 software. For normally distributed data, parametric tests were used to compare mean differences between two groups and results were analyzed by unpaired student’s t-test. To determine mean differences between three or more groups the Two-way ANOVA with repeated measures **(RM)** were used depending on the number of factors and level of factors (ie., within-subjects, between­ subjects) to be analyzed. If the main effects between groups or within-subjects reach statistical significance (*p<0.05), an appropriate post-hoc analysis (i.e., Bonferroni) was performed.

## Results

### Chronic MEHP exposure does not affect spatial learning, recognition and/or short-term memory

To assess whether chronic MEHP administration impacts spatial learning and memory, the Y-maze test was used and spontaneous alternation behavior was quantified (Miedel et al., 2017). This test is based on an animal’s willingness to explore new arms of the maze and mice typically show a tendency to enter an arm that was recently visited. If the mouse enters a different arm for 3 consecutive arm entries, a spontaneous alternation occurs (Miedel et al., 2017). MEHP-treated mice and vehicle mice showed a similar percentage of spontaneous alternations (**Figure 2A-B** *Vehicle* 53.97± 5.14 vs *MEHP* 52.21± 3.91, *p*>0.05), suggesting the MEHP exposure did not impair innate exploration and spatial memory.

**Figure 2:**
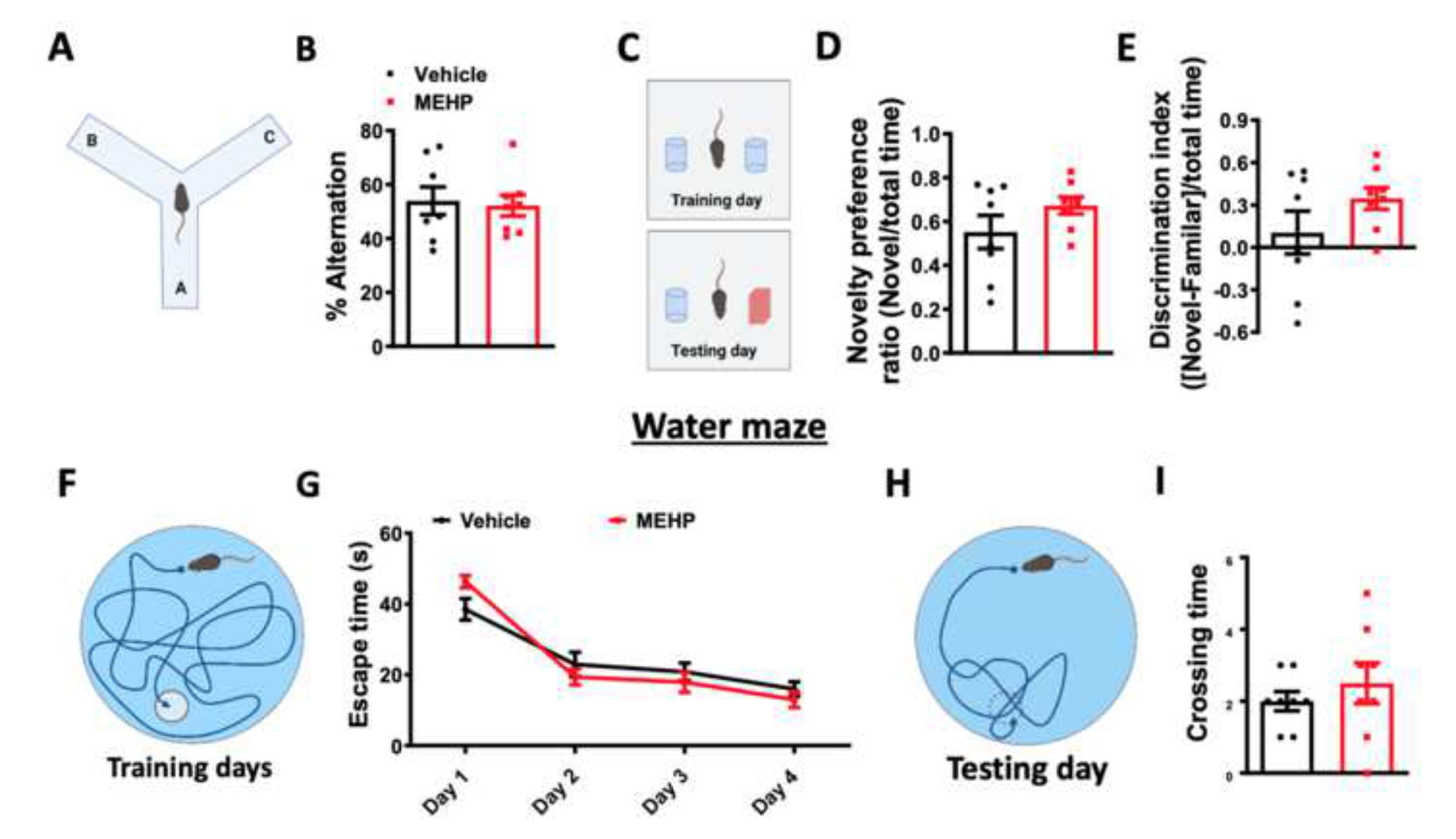
The effects of chronic MEHP on spatial learning, recognition and/or short-term memory: (A) Experimental protocol for Y-maze test. (B) Percentage of spontaneous alternation in the Y-maze test. (C) Experimental protocol for novel object recognition test. (D) Novelty preference ratio in the novel object recognition test. (E) Discrimination index in the novel object recognition test. (F) Experimental protocol for water maze training days. (G) Escaping time in the water maze training. (H) Experimental protocol for water maze testing. (I) Time that the mice crossed the position where the platform used to be during water maze testing. Data are presented as mean± SEM, n = 8.

Rodents have an innate preference for novelty and the choice to explore the novel object reflects the use of learning and recognition memory (Leger et al., 2013). Therefore we next evaluated the effects of MEHP on cognition or recognition memory, using the novel object recognition test (**Figure 2C-E**). Mice were allowed to explore an open field containing two objects, and on the second day, one of the objects was replaced with a novel one. As shown in **Figure 2C**, differences in the exploration time of novel and familiar objects were used to evaluate learning and memory. Both groups of mice showed similar novelty performance (novel object exploring time/ total exploring time, *Vehicle* 0.55± 0.08 vs *MEHP* 0.67±0.04, *p*>0.05) and discrimination index ((novel object exploring time - familiar object exploring time)/ total exploring time, *Vehicle* 0.10± 0.15 vs *MEHP* 0.35±0.08, *p*>0.05) (**Figure 2D-E**).

Finally, to assess the effects of MEHP on long-term spatial memory, a Morris water maze (MWM) was conducted. As shown in **Figure 2F and H**, mice were introduced to the water maze and the amount of time spent searching for a hidden platform (escape time) was recorded. On the last day of the test, the platform was removed, and the number of times that mice crossed the position where the platform was previously located were measured (crossing time). No significant differences were found in escape time (F_(1,14)_ = 0.024, *p*>0.05) or crossing time *(Vehicle* 2.0±0.27 vs *MEHP* 2.5±0.57, *p*>0.05) between the MEHP and the vehicle groups (**Figure 2G**, I), suggesting that long-term spatial memory was not impaired by MEHP exposure.

### Chronic MEHP exposure increases anxiety-like behaviors

Using elevated plus maze (EPM) and open field testing (OFT), we next assessed the effects of chronic phthalate exposure on anxiety-like behaviors. As shown in **Figure 3B**, the total distance traveled during the 5 minutes of the EPM testing was comparable between the two groups *(Vehicle* 11.16±0.60 vs *MEHP* 10.07±0.71, *p*>0.05), however, the MEHP-treated mice performed significantly less open-arm entries (**Figure 3C**, *Vehicle* 15.42±1.29 vs *MEHP* 9.25±1.15, *p*<0.01), traveled less distance in the open arms (**Figure 3D**, *Vehicle* 2.54± 0.44 vs *MEHP* 1.27± 0.31, *p*<0.05), and spent significantly less time in the open arms (**Figure 3E**, *Vehicle* 30.70±3.93 vs *MEHP* 17.24±3.25, *p*<0.05) compared to the vehicle group.

**Figure 3:**
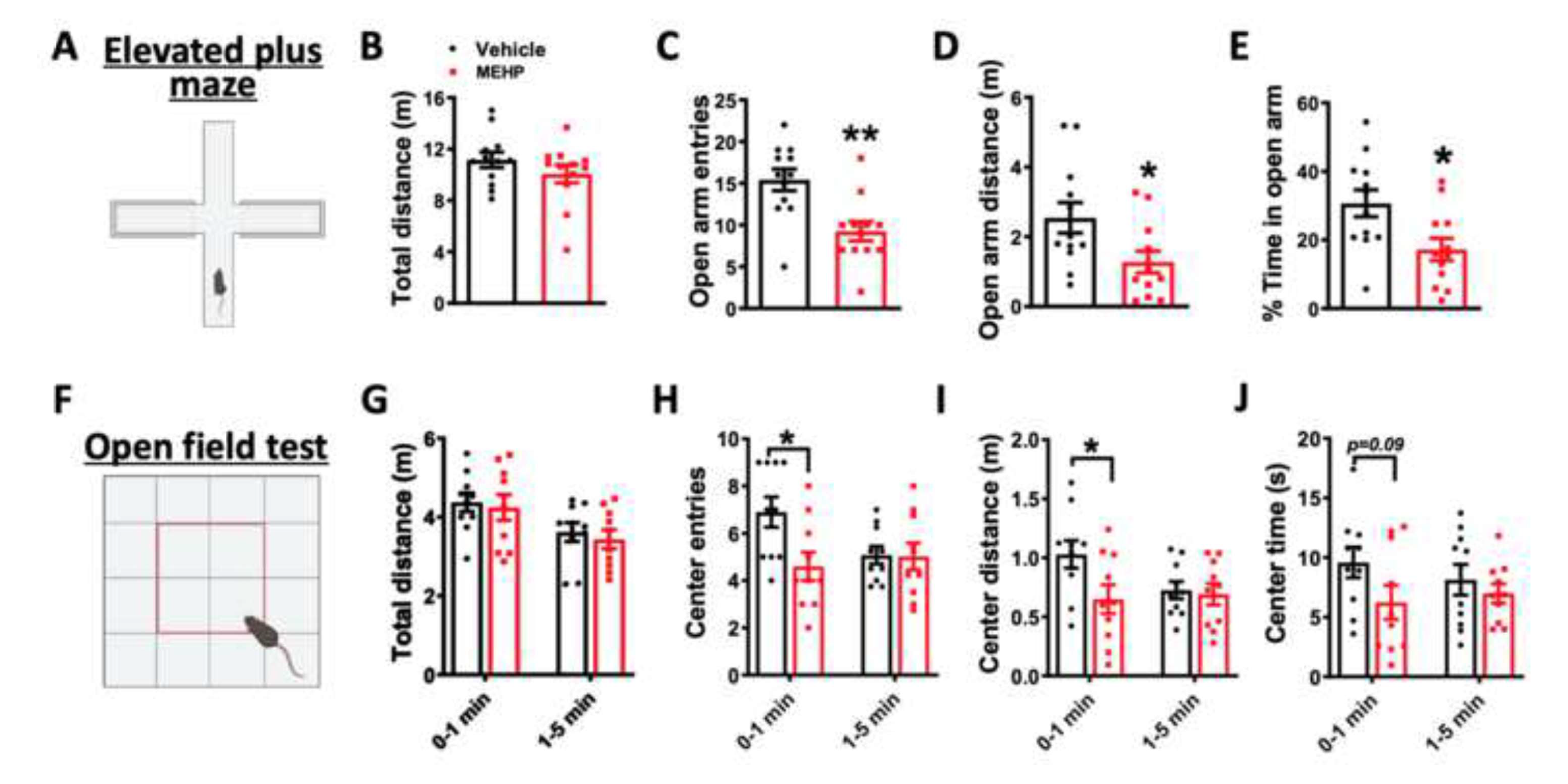
The effects of chronic MEHP on Anxiety-related measurements: (A) Experimental protocol for elevated plus maze (EPM) test. (B) Total distance traveled in EPM. (C) Open arm entries in EPM. (D) Distance traveled in the open arm of the EPM. (E) Percentage time spend in the open arm during the EPM test. (F) Experimental protocol for open field test. (G) Total distance traveled in the open field. (H) Time of entries into the center zone of the open field test. (I) Distance traveled in the center zone of the open field test. (J) Time spent in the center zone of the open field test. Data are presented as mean± SEM. *, p<0.05, **, p<0.01, two tailed Student’s t-test, n = 10-12.

To further assess the anxiolytic behavioral effects of MEHP exposure (**Figure 3 C-E**), an open field test was employed with center exploration used as an index of anxiety level. Within the first minute of testing, MEHP-treated animals had significantly fewer center entries an traveled less center distance (**Figure 3H-I** *center entries: Vehicle* 6.9± 0.64 vs *MEHP* 4.6± 0.6, *p<0.05; center distance: Vehicle* 1.03± 0.12 vs *MEHP* 0.65± 0.12, *p*<0.05), but thereafter, these parameters became comparable between MEHP-treated and control mice during the 5 min test (**Figure 3H-J** *center entries:, Vehicle* 5.08± 0.37 vs *MEHP* 5.03± 0.57, *p*>0.05; *center distance: Vehicle* 0.73± 0.07 vs *MEHP* 0.69± *0.09, p>0.05).* The increased anxiety-like behavior that accompanied MEHP administration was not a result of locomotor impairment, since the total distance traveled in the open field remained unchanged between the two groups (**Figure 3G** *0-1 min: Vehicle* 4.38± 0.24 vs *MEHP* 4.24± *0.33, p>0.05; 1-5 min: Vehicle* 3.62± 0.24 vs *MEHP* 3.44± 0.24, *p*>0.05). Locomotor activity was also evaluated for 30 min in the open field and no differences were found between the two groups (data not shown).

### Chronic MEHP exposure increases fear-related and startle reactivity

To assess the effect of MEHP exposure on learned fear and startle reactivity, an acoustic startle test and a conditioned fear test were performed (**Figure 4A, D and G**). Compared to the vehicle control, MEHP-treated mice exhibited a higher startle magnitude to the 120 dB white noise bursts, but only during the initial stage of the test (**Figure 4B**, interaction of group x burst, F (15, 330) = 2.197, *p*<0.01). Furthermore, fear responses as measured by percentage freezing to the conditioned tones (CS) were also increased in the MEHP exposed mice. As shown in **Figure 4E-F and H-l**, increased conditioned freezing responses were found in the late stage of fear acquisition (**Figure 4F**, CS­ *US 4-5: Vehicle* 64.56±3.36 vs *MEHP* 76.08±3.55, *p*<0.05). While this enhanced CS-dependent freezing in the **MEHP** group carried over to the next day during fear expression and extinction testing (**Figure 41**, *Vehicle* 56.75±3.89 vs *MEHP* 67.98±3.6, *p*<0.05).

**Figure 4:**
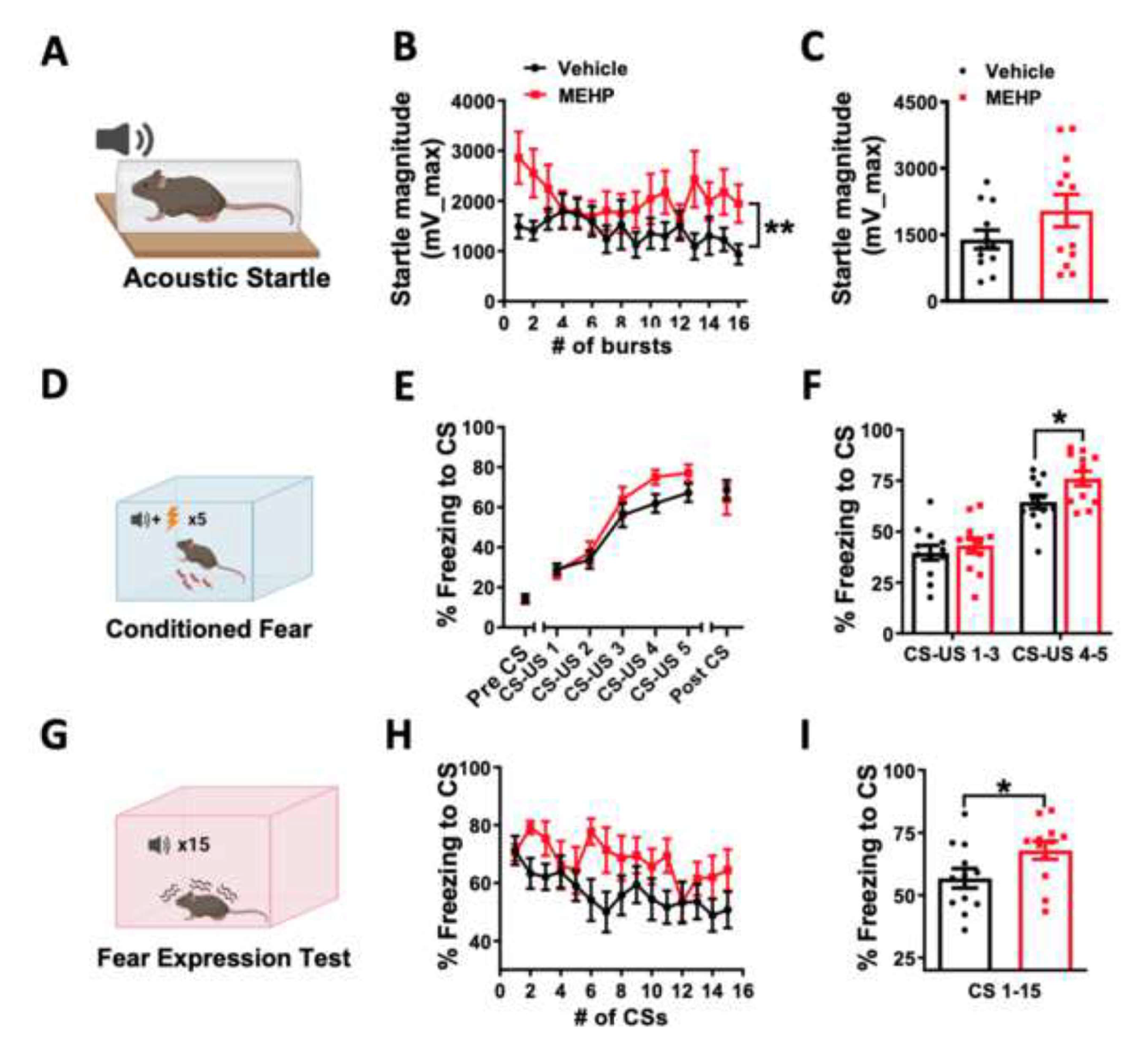
The effects of chronic MEHP on the acoustic startle response and conditioned fear: (A) Experimental protocol for acoustic startle test. (B) Startle magnitude to 120 dB white noise. Each data point represents one acoustic burst. (C) Average startle magnitude across the 16 acoustic bursts. **, significant groups x burst interaction, p<0.01, 2-Way ANOVA. (D) Experimental protocol for conditioned fear learning. (E) Percentage freezing during fear learning. (F) Average freezing percentages of the CS-US 1-3 and CS-US 4-5 during fear learning*, p<0.05, two tailed Student’s t-test. (G) Experimental protocol for fear testing. (H) Percentage freezing during fear testing. (I) Average freezing percentages during fear testing. *p<0.05, two tailed Student’s t-test. n = 12 mice /group. CS, conditioned stimulus. US, unconditioned stimulus.

### The effects of MEHP exposure on neuroendocrine and stress biomarkers

Anxiety disorders have been linked to a dysregulation of the neuroendocrine system, while phthalates can also act to disrupt the endocrine system as they are well known endocrine-disrupting chemicals. Therefore we next evaluated the effects of MEHP exposure on changes in plasma corticosterone, adrenaline, noradrenaline, dopamine concentrations and inflammatory cytokines. As shown in Figure *SA,* MEHP administration significantly increased plasma corticosterone levels *(Vehicle* 67.72±14.21 vs *MEHP* 154.5±16.40, *p*<0.01), while the concentration of adrenaline *(Vehicle* 5.89±0.87 vs *MEHP* 6.17±0.44, *p*>0.05), noradrenaline *(Vehicle* 15.12±2.72 vs *MEHP* 11.26±0.8, *p>0* .05), and dopamine *(Vehicle* 1.83±0.23 vs *MEHP* 1.71±0.11, *p>0*.05) remained unchanged (**Figure 5B-D**). Furthermore, several epidemiological and experimental animal studies have demonstrated a strong association between phthalate exposure and inflammation (Duan et al., 2017; Trim et al., 2021; Win-Shwe et al., 2013). Supporting this, MEHP-treated animals exhibited a significant increase in circulating pro-inflammatory cytokine levels **(Figure 5E-H).** including elevated IFN-γ *(Vehicle* 0.3±0.04 vs *MEHP* 0.56±0.11, *p*<0.05) and IL-2 *(Vehicle* 0.61±0.04 vs *MEHP* 1.12±0.12, *p*<0.01). Changes in inflammatory biomarkers and immune function can be a cause or consequence of endocrine or glucocorticoid disturbances and therefore we next evaluated the effects of MEHP on glucocorticoids and HPA-axis function.

**Figure 5:**
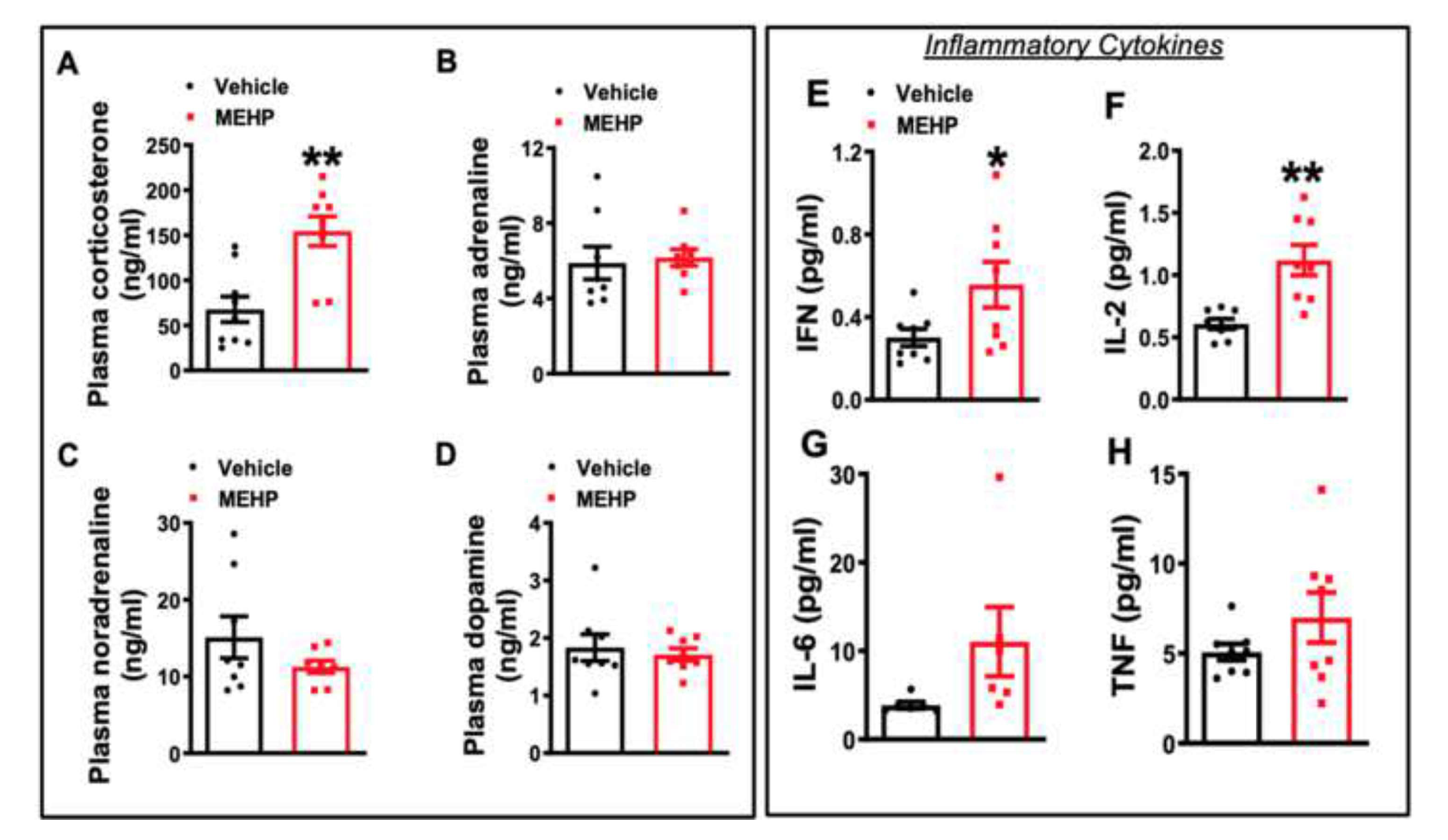
The effects of chronic MEHP on plasma corticosterone, 3-catecholamines and pro-inflammatory cytokines levels: (A) Plasma corticosterone concentration. (B) Plasma adrenaline concentration. (C) Plasma noradrenaline concentration. (D) Plasma dopamine concentration. *p<0.05, **p<0.01, two tailed Student’s t-test.

### The effects of MEHP exposure on glucocorticoid metabolic pathways

To test the hypothesis that MEHP alters HPA axis activity (**Figure 6** schematic) and consequent increases in plasma corticosterone level, we first examined mRNA levels of CRH and c-Fos (index of neuronal activity) in the hypothalamus. As shown in **Figure 6B-D**, despite a trend for a reduction in c-Fos mRNA expression, there were no statistically significant differences in hypothalamic CRH mRNA expression or plasma ACTH levels in MEHP-treated animals. These data suggest that our observed elevated plasma corticosterone levels (**Figure 5B-D**) following MEHP exposure, may not be a consequence of enhanced hypothalamic, CRH-ACTH activation but possibly as a result of downstream alterations in glucocorticoid metabolism as a result of MEHP exposure. Our next studies sought to examine this possibility.

**Figure 6:**
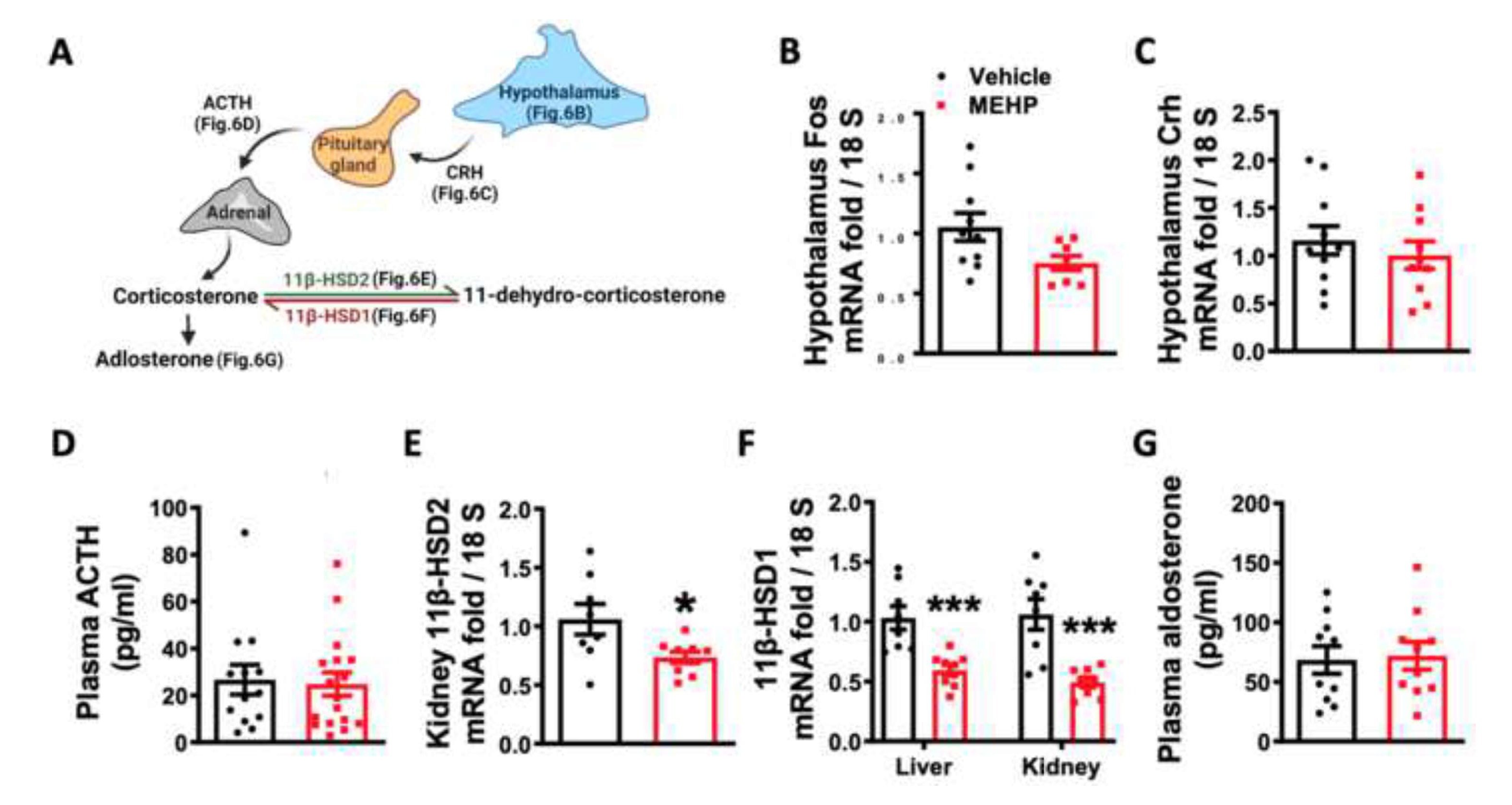
The effects of chronic MEHP on glucocorticoid metabolic pathways: Diagram illustrating steps to regulate corticosterone secretion and metabolism that were tested. (B) Neuronal activity marker c-fos mRNA expression in the hypothalamus. (C) Corticotropin-releasing hormone (CRH) mRNA expression in the hypothalamus. (D) Adrenocorticotropic hormone (ACTH) level in plasma. (E) 11β-HSD-2 mRNA expression in kidney. (F) 11β-hydroxysteroid dehydrogenase (11β-HSD)-1 mRNA expression in the liver and kidney. (F) Plasma aldosterone concentration. *p<0.05, ***p<0.001, two tailed Student’s t-test.

Corticosterone plasma concentrations, synthesis and degradation, are regulated in part by 11β­hydroxysteroid dehydrogenase (11β-HSD) enzyme activity, and the interconversion of active and inactive glucocorticoids (Chapman et al., 2013). For example, corticosterone is metabolized by 11β-HSD2 to its inactive form 11-dehydrocorticosterone, while regeneration can occur via 11β-HSDl (Chapman et al., 2013 and **Figure 6A pathway).** Accordingly, we next investigated the effects of chronic MEHP on 11β-HSD, by quantifying mRNA expression of 11β-HSDl and 11β-HSD2 in liver and kidney tissue. We hypothesized that either increased 11β-HSDl or decreased 11β-HSD2 expression could lead to increased plasma corticosterone levels in the MEHP group as a result of increased conversion of 11-dehydrocorticosterone to active corticosterone. As shown in **Figure 6E**, 11β-HSD2 was lower in kidney samples collected from MEHP-treated animals *(Vehicle* 1.06± 0.13 vs *MEHP* 0.74± *0.05, *p*<0.05),* while 11β-HSD2 expression was undetectable in liver tissue sample. MEHP-treated animals also showed decreased 11β-HSDl expression in both liver and kidney tissue (**Figure 6F**, *liver: Vehicle* 1.03± 0.1 vs *MEHP* 0.59± 0.04, *p*<0.001; *kidney: Vehicle* 1.06± 0.13 vs *MEHP* 0.49± 0.04, *p*<0.001), which may suggest a negative feedback mechanism in response to increased plasma corticosterone levels. Furthermore, changes in 11β-HSD expression can also affect the concentration of corticosterone catalytic product aldosterone, however, there we no significant differences between groups in circulating aldosterone plasma levels (**Figure 6G**, *Vehicle* 68.45± 11.56 vs *MEHP* 71.82± 11.65, *p>0* .05).

## Discussion

Due to its low cost of production and superior chemical properties, di-2-ethylhexyl phthalate (DEHP) remains the most widely used plasticizer in polyvinyl chloride (PVC) products. Once in the body, DEHP is rapidly hydrolyzed to mono-2-ethylhexyl phthalate (MEHP), which has been shown to exhibit toxicological effects on multiple organ systems - including the liver, heart, testes, and pituitary gland (Hong et al., 2009; Ito et al., 2019; Jaimes et al., 2019). Moreover, emerging studies have documented associations between prenatal or early childhood phthalate exposure on neuropsychiatric and adverse behavioral outcomes (Ejaredar et al., 2015; Zhang et al., 2019), however the effects of phthalates on postnatal behavioral outcomes are less studied. Using an experimental adult rodent model, we aimed to examine the effects of MEHP exposure on cognition, emotion and anxiety-like behaviors and their impact on neuroendocrine and inflammatory function. Our results show that MEHP exposure induces anxiety-like behavior, including enhanced fear and startle reactivity, but does not affect short-or long-term memory in adult mice. We also observed that MEHP exposure increases plasma corticosterone levels and decreases peripheral tissue mRNA expression of 11β-HSD2, which may lead to changes in neuroendocrine and immune homeostasis contributing to enhanced anxiety-like behaviors.

Growing evidence links chronic phthalate exposure to neurobehavioral outcomes (Huang et al., 2019; Lee et al., 2016) and phthalate exposure can precipitate anxiety-like behavior in rodents (Carbone et al., 2019; Hatcher et al., 2019; Ma et al., 2015; Park et al., 2015). For example, mice exposed in utero to DEHP displayed fewer social behaviors, and this effect was transgenerational and persisted to the F3 generation (Quinnies et al., 2015, 2017). Depression and impaired cognitive deficits were also reported in DEHP or DINP exposed mice (Ma et al., 2015; Wang et al., 2014; Xu et al., 2015). In the current study, we found that exposure to MEHP during a post-developmental adult period similarly drives abnormal neurobehavioral changes, which likely involve distinct temporally linked biological mechanisms that are unique from pre­ natal, gestational or transgenerational phthalate exposure, as shown in previous studies(Lovekamp-Swan & Davis, 2003; Sun et al., 2022; Tetz et al., 2015). In our study, MEHP-exposed adult mice exhibited increased anxiety-like behaviors, with evidence pointing to an enhanced systemic neuroendocrine and inflammatory response since 3-week MEHP exposure produced a significant basal increase in corticosterone levels and circulating pro-inflammatory cytokines. Interestingly, this effect was independent of increases in other peripheral stress neuroendocrine biomarkers (ie., catecholamines, aldosterone).

In addition, to the MEHP-induced anxiogenic affects observed in the present study, as determined by OFT and EPM testing, MEHP-exposed mice also exhibited increased freezing behavior during conditioned fear learning, as well as increased basal startle reflexes to acoustic bursts of white noise. Conditioned learned fear can also be a way to assess anxiety-like behaviors in mice, by assessing physical manifestations of anxiety that can drive physiological responses to fear or threats (e.g., freezing, blood pressure, heart rate, respiration) (Swiercz et al., 2018; Turley et al., 2021). For example, mouse lines characterized by high innate anxiety-related behavior exhibit more freezing behavior in response to conditioned stimuli (Maren & Holmes, 2016; Sartori et al., 2011), indicating that stress-induced anxiety enhances the physiological expression of conditioned fear. Furthermore, the startle reflex is also linked to anxiety, as individuals with anxiety disorders show a greater startle reflex (Poli & Angrilli, 2015; Ray et al., 2009). The elevated freezing and startle responses observed in the MEHP mice, thus support an anxiogenic phenotype and enhanced conditioned fear-related cardiovascular reactivity (Jaimes et al., 2017). Despite the MEHP induced anxiogenic affects, we did not observe deficits in measures of cognition, short or long-term memory testing following MEHP, thus suggesting an MEHP dependent neurobehavioral anxiogenic phenotype, independent of changes in measures of cognition and memory. These results contrast with some studies demonstrating phthalate induced impairments in rodent learning and memory. However in these experiments phthalate administration was given pre or perinatally, suggesting a developmental phthalate affect on cognition, as opposed to the current study where MEHP was administered in adult mice.

Neuroendocrine biomarker or hypothalamus-pituitary-adrenal (HPA)-axis dysregulation is a common corollary of anxiety and depressive phenotypes (Elsaafien et al., 2021; Herman et al., 2016; Mitra & Sapolsky, 2008). Phthalates are known neuroendocrine disruptors and therefore can alter the homeostasis of the neuroendocrine system and have significant detrimental neurodevelopmental effects on central nervous system function, specifically, the HPA-axis. To assess this, following MEHP exposure, we first measured circulating levels of corticosterone and its pre-cursor adrenocorticotropic hormone (ACTH) (Figure 6).

Following 3 weeks of MEHP administration, basal corticosterone levels were significantly elevated, with an absence of a change in an expected corresponding increase in ACTH (Herman et al., 2016). The dissociation between ACTH levels and corticosterone release is likely due to lag times in the temporal dynamics of glucocorticoid feedback signaling and degradation processes (Herman et al., 2016). Similarly, at the hypothalamic level tissue level, chronic MEHP treatment did not alter HPA-axis activity, as indicated by an absence of an increase in hypothalamus CRH mRNA expression. However, there was a trend for a reduction in the activity of hypothalamus neurons *(c-Fos* marker) after MEHP administration, which may be a result of increased negative hypothalamic feedback in response to plasma corticosterone levels. Overall, our neuroendocrine results suggest that MEHP-dependent increases in corticosterone are likely a consequence of activity outside the pituitary at the level of the adrenal gland or other peripheral tissues (e.g., kidney, liver) that regulate peripheral glucocorticoid metabolism and circulating corticosterone levels (**Fig. 6A**).

For example, previous studies has shown that phthalates can affect glucocorticoid activity by decreasing RNA levels and enzyme activity of the 11- hydroxysteroid dehydrogenase 11- 2 (HSD2), which in tum converts corticosterone into its receptor-inactive form 11- dehydrocorticosterone (Vitku et al., 2016). Clinical data suggest increased phthalate exposure is associated with elevated cortisol/cortisone ratio in premature infants, which may be mechanistically linked to the inhibition of 11- HSD2 (Jenkins et al., 2019). Our current results indicate that chronic MEHP administration decreases mRNA levels of 11- HSD2, which could contribute to increased circulating levels of corticosterone. However, contrary to what we expected, the mRNA levels of the type 1 isoform-11- HSDl, were decreased after MEHP administration. Overall, we speculate that the elevated corticosterone levels following chronic MEHP exposure maybe due in part to altered negative feedback as a consequence of altered HSD enzymatic function and glucocorticoid metabolism.

Glucocorticoid dysregulation is also tightly linked to immune homeostasis and inflammation. For example, the interaction, coordination, and crosstalk between the endocrine system via the glucocorticoid receptor (GR) and immune system regulating transcription factor nuclear factor kappa-light chain-enhancer of activated B cells (NFxB), is well known for its central role in maintaining this neuroendocrine-immune balance. A scenario of high corticosterone levels, may contribute to decreased GR sensitivity and contribute to increased levels of pro-inflammatory signals and cytokines (Bekhbat et al., 2017) and contribute to the overall MEHP dependent anxiogenic phenotype, independent of any cognitive behavioral affects. Indeed, previous studies have shown that several pro-inflammatory biomarkers, are elevated following phthalate exposure (Manteiga & Lee, 2017; Murphy et al., 2014; Voss et al., 2018). In addition, various *in vitro* studies have reported that acute phthalate exposure increases the production and secretion of inflammatory cytokines from macrophages and neutrophils (e.g., TNF-a, IL-lb, IL-8, and IL-6) (Bolling et al., 2012; Mohammadi & Ashari, 2021; Nishioka et al., 2012; Vetrano et al., 2010). Although less is known about the *in vivo* effects of phthalate exposure on systemic inflammation, studies suggest that phthalates amplify proinflammatory cytokine production and increase immune cell infiltration to both testicular and cardiac tissue (Schwendt et al., 2022; Voss et al., 2018). Supporting this, we provide evidence for an increased proinflammatory state in mice chronically exposed to MEHP, which maybe a consequence of phthalate-induced cortisol elevations and a driver of anxiety-like behavior observed here. However additional studies are required to establish this mechanistic link.

In summary, chronic MEHP exposure induces anxiety-like behaviors in adult rodents without altering cognition or memory, which maybe in part due to altered HPA-axis function and peripheral glucocorticoid metabolism. These results expand the current understanding for the effects of MEHP as an endocrine disrupting and toxic environmental chemical on neurobehavioral function in adult mice. Future studies are needed to further investigate the link between phthalate exposure in adult mice, glucocorticoid regulation, and the downstream cellular, molecular and physiological pathways. Finally, given that these studies were conducted in male mice, future studies are needed to further examine the effects of MEHP exposure on the sexual dimorphisms (Adam & Mhaouty-Kodja, 2022) of endocrine and immune homeostasis.

## ACKNOWLEDGMENTS

**GRANTS:** This work was supported by the National Institutes of Health (R01HL139472 to NGP and PJM) and NIH 1R01HL137103-01Al (PJM) and (CDMRP) PR210574 (PJM).

**DISCLOSURES:** No conflicts of interest, financial or otherwise, are declared.

